# Nonclassical monocytes are prone to migrate into tumor in diffuse large B-cell lymphoma

**DOI:** 10.1101/2021.04.10.439292

**Authors:** Simon Le Gallou, Faustine Lhomme, Jonathan M. Irish, Anna Mingam, Celine Pangault, Celine Monvoisin, Juliette Ferrant, Imane Azzaoui, Delphine Rossille, Krimo Bouabdallah, Gandhi Damaj, Guillaume Cartron, Pascal Godmer, Steven Le Gouill, René-Olivier Casasnovas, Thierry Jo Molina, Roch Houot, Thierry Lamy, Karin Tarte, Thierry Fest, Mikael Roussel

## Abstract

Absolute count of circulating monocytes has been proposed as an independent prognostic factor in diffuse large B-cell lymphoma (DLBCL). However, monocyte nomenclature includes various subsets with pro-, anti-inflammatory, or suppressive functions, and their clinical relevance in DLBCL has been poorly explored. Herein, we broadly assessed circulating monocyte heterogeneity in 91 DLBCL patients. Classical- (cMO, CD14^pos^ CD16^neg^) and intermediate- (iMO, CD14^pos^ CD16^pos^) monocytes accumulated in DLBCL peripheral blood and exhibited an inflammatory phenotype. On the opposite, nonclassical monocytes (ncMO, CD14^low^ CD16^pos^) were decreased in peripheral blood. Tumor-conditioned monocytes presented similarities with ncMO phenotype from DLBCL and were prone to migrate in response to CCL3, CCL5, and CXCL12, and presented similarities with DLBCL-infiltrated myeloid cells, as defined by mass cytometry. Finally, we demonstrated the adverse value of an accumulation of nonclassical monocytes in 2 independent cohorts of DLBCL.

**Key points:**

- Nonclassical monocytes are prone to migrate to DLBCL tumor
- High count of circulating nonclassical monocytes is an independent adverse event in DLBCL

## Introduction

Circulating monocytes are classified by their CD14 and CD16 expression as classical- (cMO, CD14^pos^ CD16^neg^), intermediate- (iMO, CD14^pos^ CD16^pos^), and nonclassical- monocytes (ncMO, CD14^low^ CD16^pos^).^1^ In addition, Slan expression (6-Sulfo LacNac, which is a carbohydrate modification of P-selectin glycoprotein ligand-1 [PSGL-1]), allows the sub-classification of ncMO Slan^pos^ (CD14^low^ CD16^pos^ Slan^pos^) and ncMO Slan^neg^ (CD14^low^ CD16^pos^ Slan^neg^).^2,3^ Lastly monocytic myeloid derived suppressor cells (M-MDSC, CD14^pos^ HLA-DR^low^) found in acute or chronic inflammatory context, including cancers, are defined by an impairment of T- and NK- effector functions.^4^ This nomenclature reflects pro-inflammatory, anti-inflammatory, or suppressive functions described for monocytes.^5,6^

In diffuse large B-cell lymphoma (DLBCL), tumor microenvironment (TME), myeloid cells are supportive of the neoplastic process.^7–10^ In blood from DLBCL patients, an increase in circulating monocytes is a marker of adverse prognosis.^11–15^. However, so far monocytes were considered as a whole, and few studies analyzed the monocyte subsets and their clinical relevance even if their intrinsic functions are known to be different. Among monocyte subsets: i) Slan^pos^ monocytes were increased and displayed high rituximab mediated antibody-dependent cellular cytotoxicity;^16^ ii) an increase in CD16^pos^ or CD11b^pos^CX3CR1^pos^ monocytes predicted poor progression free- and overall- survival;^17,18^ iii) CD14^pos^CD163^pos^PD-L1^pos^ monocytes were increased;^19^ and finally iv) functional M-MDSCs were enriched in peripheral blood and predicted poor event-free survival.^20–22^ In DLBCL tumor, the myeloid compartment heterogeneity was recently approached by high dimensional analysis revealing distinct macrophage phenotype across lymphoma subtypes.^23^

In light with the observation that various monocyte subsets are involved in the biology of DLBCL, we investigated the canonical cMO, iMO, ncMO Slan^pos^, and ncMO Slan^neg^ subsets in two large cohorts of patients. We quantified these subsets, analyzed their phenotype and functions as well as the clinical relevance of these cells. We found here that in DLBCL, ncMO are prone to migrate into tissues and that their increase in peripheral blood is associated with an adverse prognosis.

## Methods

### Samples

A cohort of 91 DLBCL patients at diagnosis from the BMS-LyTRANS clinical trial (ClinicalTrials.gov Identifier: NCT01287923) was used in this study. Clinical characteristics of DLBCL patients enrolled in this training cohort are listed in Table 1. Patients with previous corticosteroid treatment were excluded from this study. As controls, age-matched heathy donors (HD, n = 49), follicular lymphomas (n = 9), mantle cell lymphomas (n = 9), chronic lymphocytic leukemias (n = 11), and marginal zone lymphomas (n = 10) were included. Part of these samples (DLBCL and HD) were used in a previous work.^22^ Prognosis scores were validated in a second cohort of 155 DLBCL patients from the recently published GAINED trial (ClinicalTrials.gov Identifier: NCT01659099).^24^ Clinical characteristics of DLBCL patients enrolled in this validation cohort are listed in Table 1. Finally, we reanalyzed CyTOF data (Flow Repository FR-FCM-Z2CA, already published by our group)^23^ from myeloid cells from DLBCL tumors (n = 7). The research protocol was conducted under French legal guidelines and fulfilled the requirements of the local institutional ethics committee.

**Table 1:** Patient’s characteristics at baseline.

|  | Healthy donors<br>(n = 49) | BMS-LyTRANS<br>Training cohort<br>(n = 91) | GAINED<sup>24</sup><br>Validation cohort<br>(n=155) |
| --- | --- | --- | --- |
| **Average age, years (range)** | 52.9 (25-66) | 55.7 (18-83) | 44.9 (19-60) |
| **Male, n (%)** | 30 (61.2%) | 56 (61.5%) | 86 (55.5%) |
| **Female, n (%)** | 19 (38.8%) | 35 (38.5%) | 69 (44.5%) |
| **aalPI, n (%)** |  |  |  |
| 0 to 1 | NA | 46 (59%)* | 68 (43.9%) |
| 2 | NA | 28 (35.9%)* | 71 (45.8%) |
| 3 | NA | 4 (5.1%)* | 16 (10.3%) |
| **Cell of origin, n (%)** |  |  |  |
| GCB (Hans<sup>&</sup> or Nanostring<sup>@</sup>) | NA | 30 (54.5%)*,<sup>&</sup> | 83 (74.8%)*,<sup>@</sup> |
| Non-GCB (Hans<sup>&</sup>) or ABC (Nanostring<sup>@</sup>) | NA | 25 (45.5%)*,<sup>&</sup> | 28 (25.2%)*,<sup>@</sup> |
| Unclassified<sup>@</sup> | NA | NA | 9<sup>@</sup> |
| Insufficient material | NA | 36 | 35 |
| **Chemotherapy, n** |  |  |  |
| Rituximab CHOP | NA | 69 | 37 |
| Rituximab ACVBP | NA | 0 | 39 |
| Obinutuzumab CHOP | NA | 6 | 40 |
| Obinutuzumab ACVBP | NA | 0 | 38 |
| Other treatment | NA | 3 | 0 |
| Unknown or not treated | NA | 13 | 1 |

aalPI: age adjusted International prognostic index; GCB: germinal center B cell; ABC: activated B cell \*: percentage among cases with known data; &: defined by Hans algorithm; @: defined by nanostring analysis. CHOP: cyclophosphamide, doxorubicine, vincristine, and prednisone; ACVBP: doxorubicine, prednisone, cyclophosphamide, vindesine, and bleomycine; NA: not applicable.

### Fluorescent flow cytometry analysis

Blood samples were collected on heparin tubes. Flow cytometry analysis of M-MDSCs, cMO, iMO, and ncMO were performed on whole blood (300 μL/tube) with the antibody panel shown Table S1 and the gating strategy defined Figure S1. Absolute counts were obtained by using Flow-Count beads (Beckman Coulter, Brea, CA). An erythrocytes lysis (Uti-Lyse Dako, Carpinteria, CA) was performed before analysis by flow cytometry (Navios, Beckman Coulter). Analyses were performed using Kaluza software (Beckman Coulter).

### In vitro culture

Monocytes were obtained from PBMCs by elutriation before cryopreservation (plate-forme DTC; CIC Biotherapie, Nantes, France). Monocytes were thawed and resuspended at 4 x10^6^ cells/mL in RPMI 1640 (Invitrogen, Carlsband, CA, USA) supplemented with 10 % FCS and antibiotics (Invitrogen) and then diluted at 2 x10^6^ cells/mL by adding, as control, the OCI-Ly medium (IMDM supplemented with 10% human AB serum,1% penicillin–streptomycin and 50 μM of β-Mercaptoethanol, or OCI-Ly3 or OCI-Ly19 supernatant. Two mL of cell suspension were seeded in a 6-well plate during 4 days before mass cytometry analysis or migration assay.

### Mass cytometry analysis

Cell labeling and mass-cytometry analysis were performed as previously described.^25–27^ Briefly, cells were incubated with 25 μM cisplatin (Fluidigm San Francisco, CA, USA). Then, 5 x10^6^ cells were washed in PBS (HyClone Laboratories, Logan, UT, USA) containing 1 % BSA (Thermo Fisher Scientific) and stained in 100 μL PBS and BSA 1 %-containing Antibody cocktail. Cells were stained for 30 min at RT with the antibodies (Table S2) Cells were washed twice in PBS - BSA 1 % before fixation in 1.6 % PFA, and permeabilization with methanol (Electron Microscopy Sciences, Hatfield, PA, USA). After incubating overnight at −20°C in MeOH, cells were washed twice with PBS -BSA 1 % and stained 20 min with iridium intercalator (Fluidigm, Sunnyvale, CA, US). Finally, cells were washed twice with PBS and twice with diH2O before acquisition a CyTOF 2.0 mass cytometer (Fluidigm).

### Data processing and analysis

Data analysis was performed using the workflow previously developed.^23^ Briefly, after acquisition, intrafile signal drift was normalized and .fcs files were obtained using CyTOF software. To diminish batch effects, all files were normalized on EQ Beads (Fluidigm) using the premessa R package (https://github.com/ParkerICI/premessa). Raw median intensity values were transformed to a hyperbolic arcsine (arcsinh) scale with a cofactor of 5, then analysis was performed using Cytobank software (Beckman Coulter, Brea, CA, USA). Each file was pre-gated for single, viable cells. These populations were exported as separate flow cytometry standard files and analyzed using Cyclospore.^28,29^ The HSNE (Hierarchical Stochastic Neighbor Embedding) was performed to identify cell types. Then, hierarchical clustering of mean marker intensity on each cluster representing a phenotypically distinct myeloid cell population was performed.

### Migration assay

At day 4 of monocyte culture with OCI-Ly3, or OCI-Ly19 supernatant, or control culture medium, cells were collected and washed twice in PBS before starvation during 1 hour (37°C) at 10^6^ cells/mL in RPMI 1% HSA. Cells were washed once and 100 μL of cells at 10^6^ cells/mL were added to the upper compartment of Transwell chambers with 5 μM pore filters. The lower chamber contained CCL2 (R&D Systems, 30 ng/ml), CCL3 (R&D Systems, 20 ng/ml), CCL5 (R&D Systems, 30 ng/ml), CCL22 (R&D Systems, 20 ng/ml), CXCL5 (R&D Systems, 20 ng/ml), CXCL12 (R&D Systems, 20 ng/ml), or RPMI 1640 1% HSA as control. Cells in the lower chamber were collected after 5h and the absolute number of viable (DAPI negative) monocytes was quantified by flow cytometry using FlowCount beads.

### Cell sorting

cMO (CD19^neg^ CD3^neg^ CD335^neg^ CD45^pos^ CD14^high^ CD16^neg^), iMO (CD19^neg^ CD3^neg^ CD335^neg^ CD45^pos^ CD14^high^ CD16^pos^), ncMO Slan^pos^ (CD19^neg^ CD3^neg^ CD335^neg^ CD45^pos^ CD14^low^ CD16^pos^ Slan^pos^), and ncMO Slan^neg^ (CD19^neg^ CD3^neg^ CD335^neg^ CD45^pos^ CD14^low^ CD16^pos^ Slan^neg^) were sorted from thawed PBMC of DLBCL patients and HD using an ARIA II (FACSAria, BD Biosciences).

### Quantitative real-time PCR

Total RNA was extracted usingNucleospin® RNA XS kit (Macherey-Nagel, Duren, Germany). cDNA was then generated using Fluidigm Reverse Transcription Master Mix (Fluidigm). The qPCR were performed in triplicate using 96.96 Dynamic Array™ IFCs and the BioMark™ HD System from Fluidigm. For each sample, the mean CT value for the gene of interest was calculated, normalized to the geometric mean value of the 2 housekeeping genes (*CDKN1B*, and *ELF1*) (Table S3), and compared to the median value obtained from the reference population (HD cMO or iMO for Figure 2, and DLBCL ncMO for Figure 3) using the 2-ddCT method. Results were expressed as the ratio of sample mean to reference mean for each gene.

**Figure 2:**
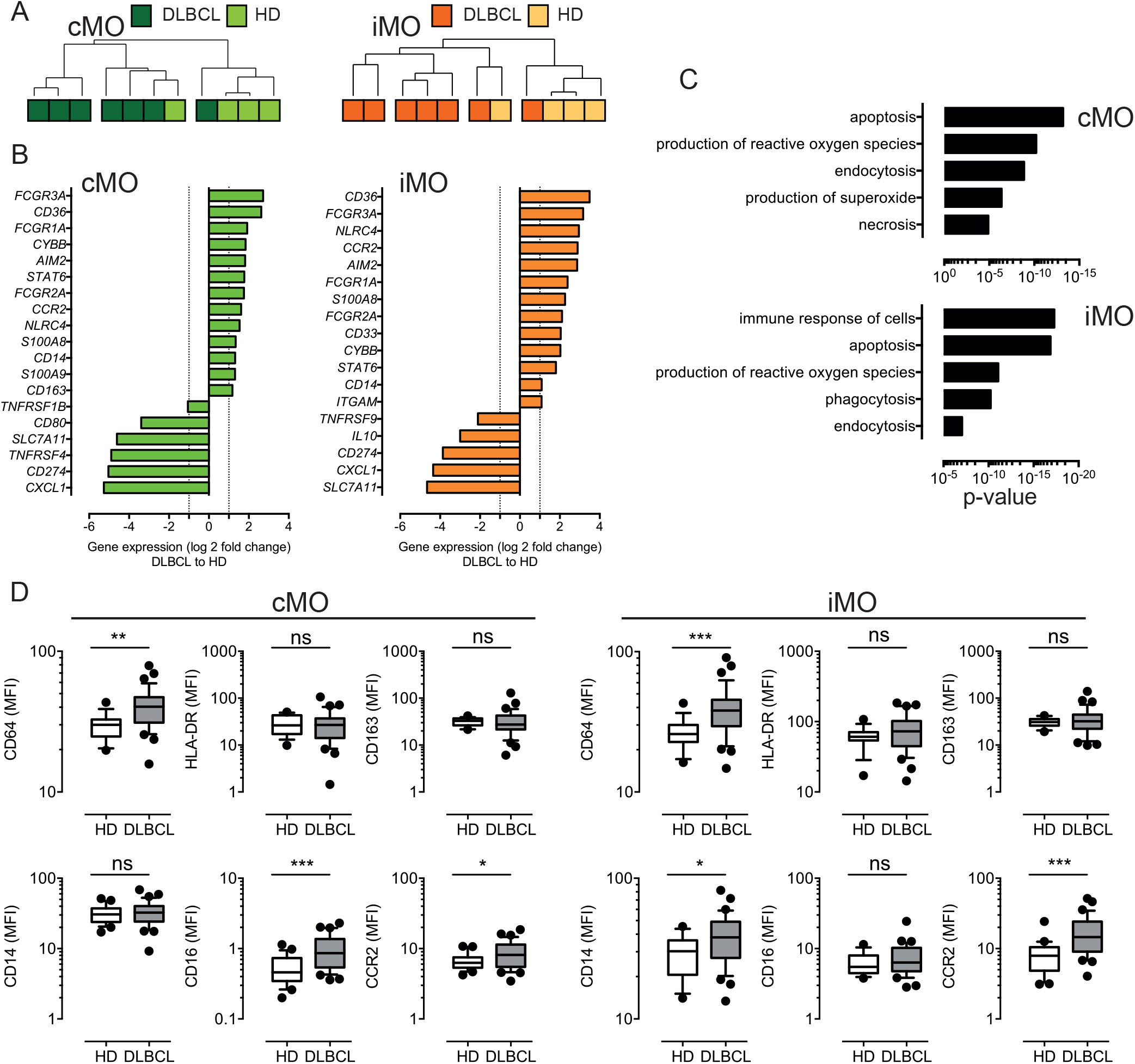
Circulating cMO and iMO share a common inflammatory phenotype, in DLBCL. (A) Hierarchical clustering of classical- (cMO) and intermediate- (iMO) monocytes, from HD (n = 4) and DLBCL (n = 7) samples. See Table S3 for a list of genes analyzed on monocyte subsets after cell sorting. Pearson’s correlation and complete linkage was employed. (B) Transcripts differentially expressed (P < .05; l1l log2FCl1l > 1) between DLBCLs and HDs, for cMO and iMO. (C) Predicted top 5 biological processes increased for cMO and iMO from DLBCL (Ingenuity Pathway Analysis, z-score > 2.5, ranked by p-value). (D) Mean fluorescence (MFI) for CD64, HLA-DR, CD163, CD14, CD16, and CCR2 for HD (n = 16) and DLBCL (n = 33) samples. **P < .01, ***P < .001, ns: non significant.

**Figure 3:**
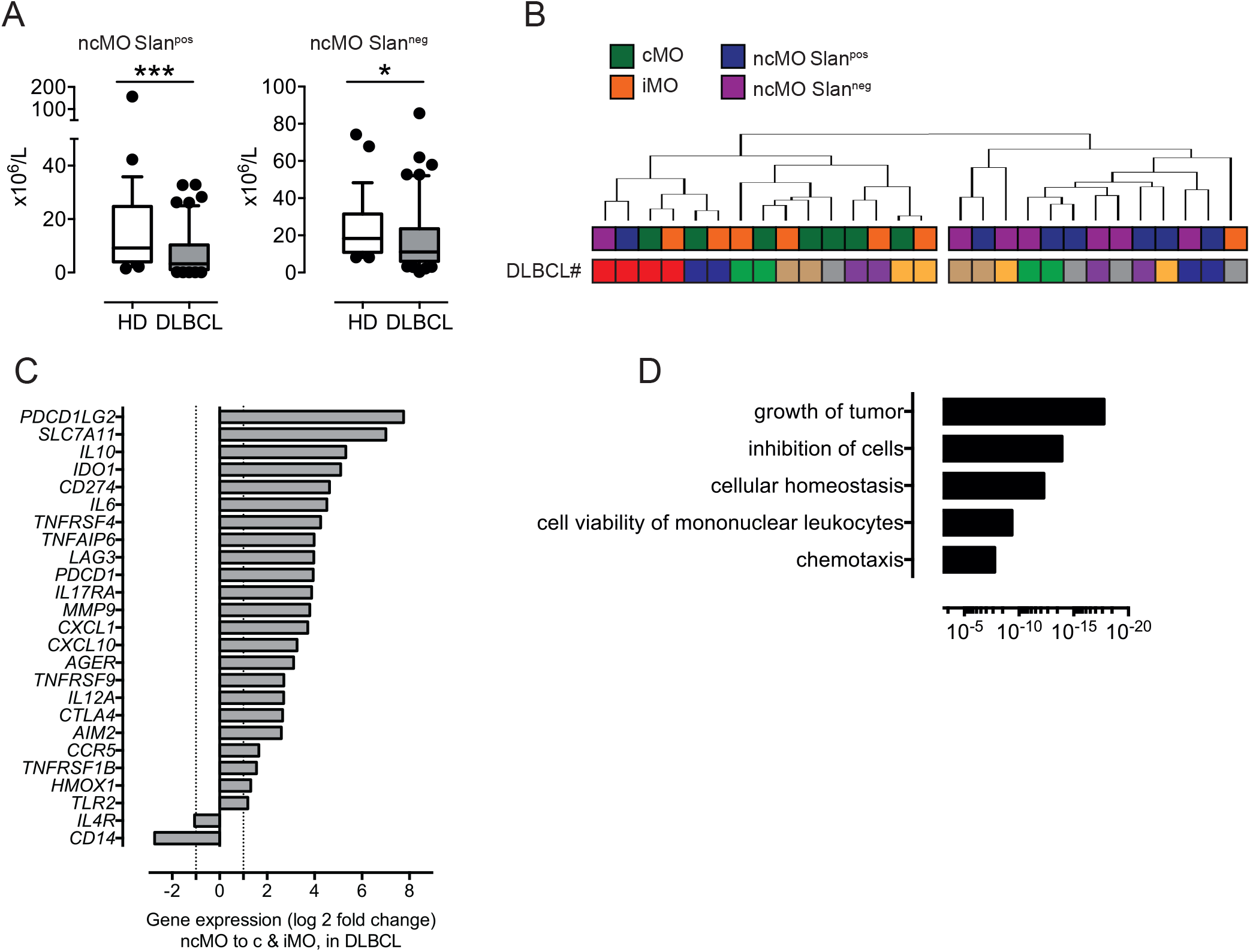
ncMO are decreased in peripheral blood but exhibit an inflammatory- and tolerogenic- phenotype. (A) Nonclassical Slan^pos^ or Slan^neg^ (ncMOSlan^pos^ and ncMoSlan^neg^) monocytes from HDs (n = 28) and DLBCLs (n = 56). (B) Hierarchical clustering of classical- (cMO), intermediate- (iMO), nonclassical Slan^pos^- or Slan^neg^- (ncMOSlan^pos^ and ncMoSlan^neg^) monocytes from DLBCL (n = 7) samples. DLBCL identity (#). List of genes analyzed on monocyte subsets after cell sorting is on Table S3. Pearson’s correlation and complete linkage was employed. (C) Transcripts (P <.05; l1l log2FCl1l > 1) enriched in ncMO compared to cMO and iMO for DLBCL. (D) Predicted top 5 biological processes increased for ncMO compared to cMO and iMO, from DLBCL (Ingenuity Pathway Analysis, z-score > 2.5, ranked by p-value).

### Statistical analysis

Statistical analyses were performed with GraphPad Prism 8.4.3 software (GraphPad Software, San Diego, CA, USA) using Spearman correlation, Wilcoxon, Mann-Whitney, Ordinary one-way ANOVA with Tukey’s multiple comparisons test, and Fishers’s exact tests as appropriate. Optimal thresholds were defined with the *maxstat* package, log-rank tests were performed with the *survmine*r package, cox model for univariate and multivariate analysis were performed with the *survival* package. Analyses were generated with R v4.0.3, using Rstudio v1.3.1093.

### Data sharing statement

For original data, please contact the corresponding author.

## Results

### cMO and iMO are increased in DLBCL

We have previously shown that M-MDSCs accumulated in DLBCL peripheral blood.^22^ However, this increase accounts for only a part of the total monocyte accumulation suggesting that additional monocyte subsets are also increased in DLBCL samples (Figure 1A). We quantified the absolute count of the 4 circulating monocyte subsets M-MDSC, cMO, iMO, and ncMO. M-MDSC, cMO, and iMO were increased in DLBCL when compared to HDs (P < .05, median: 5.75 x10^6^ cells/L vs 2.8 x10^6^ cells/L, 348.8 x10^6^ cells/L vs 274.1 x10^6^ cells/L, and 34.4 x10^6^ cells/L vs 26.1 x10^6^ cells/L; respectively). Conversely, ncMO were significantly decreased in DLBCL when compared to HD (P < .0001, median: 17.1 x10^6^ cells/L vs 36.1 x10^6^ cells/L; Figure 1B). Noteworthy, whereas the increase of cMO and iMO was also found in other B cell lymphomas, this decrease in ncMO was specific of DLBCL (Figure S2). Then, we wondered in which monocyte subset M-MDSCs were included. Of note M-MDSC count was correlated with total monocyte (R = .70, P < .0001), cMO (R = .55, P < .0001), and iMO (R = .74, P < .0001) but was not correlated with ncMO (Figure 1C) No correlation were observed between MO subsets in HD samples (data not shown). Regarding CD14 and CD16 expression, M-MDSCs were essentially aligned with the cMO phenotype and to a lesser extent to iMO (Figure 1D). Altogether, these results confirmed that in addition to MDSCs, cMO and iMO were also involved in the monocyte increase observed in DLBCL patients.

**Figure 1:**
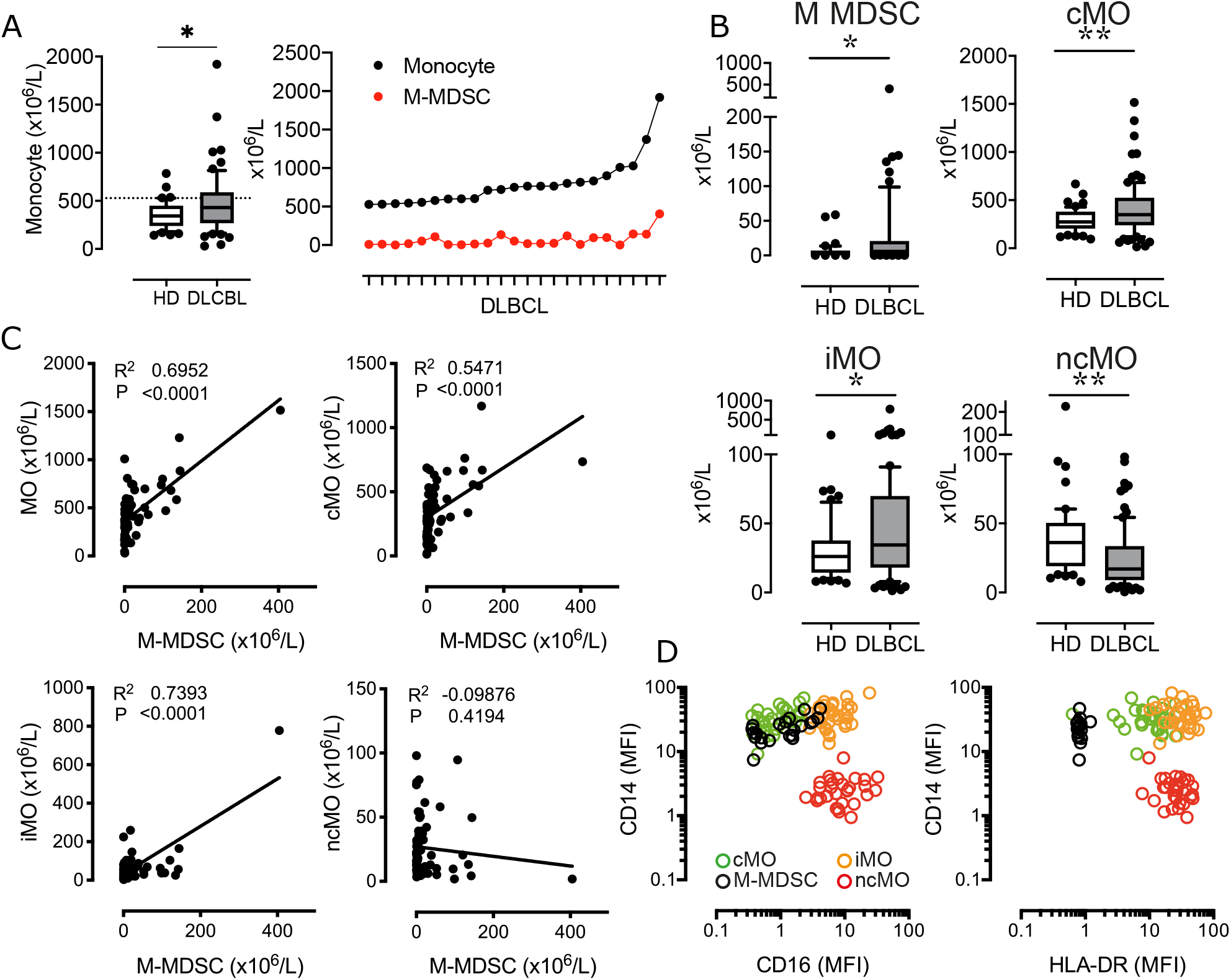
cMO and iMO are increased in peripheral blood from DLBCL. (A) Monocyte absolute counts in HD (n = 43) and DLBCL (n = 69). For the 23 DLBCL samples with monocyte above 528 x10^6^ monocytes/L (corresponding to the 90^th^ percentile of HD), proportion of M-MDSC within monocyte (B) M-MDSC in HD (n = 43) and DLBCL (n = 69) and monocyte subset counts (classical- [cMO], intermediate- [iMO], and nonclassical- [ncMO]) in peripheral blood from HD (n = 55) and DLBCL patients (n = 91). (C) Pearson correlation between M-MDSC and monocyte (MO), cMO, iMO, and ncMO (n = 69). (D)- Mean fluorescent intensity (MFI) for CD14, CD16, and HLA-DR. Each dot represents a DLBCL sample (n = 33) colored by monocyte subset (MDSC, cMO, iMO, and ncMO). *P < .05, **P < .01, ****P < .0001.

### DLBCL cMO and iMO share a common inflammatory phenotype

To further identify the immune properties of monocyte subsets, we sorted cMO and iMO from DLBCL (n=7) and HD (n=4) samples. Gene expression was assessed by high throughput qPCR on 71 genes involved in myeloid biology^22^ (Table S3 and Figure S3). Of note, 6 cMO and 5 iMO out of 7 DLBCL were clustered (Figure 2A and Figure S3). DLBCL cMO and DLBCL iMO were significantly enriched for *FCGR3A, CD36, FCGR1A, CYBB, AIM2, STAT6, FCGR2A, CCR2, NLRC4, S100A8*, and *CD14* genes, when compared to the corresponding subsets in HDs (P <.05∣log2FC∣>1) (Figure 2B). In addition, *S100A9* and *CD163* were also increased in DLBCL cMO, whereas *CD33* and *ITGAM* were enriched only in DLBCL iMO. For both subsets, *SLC7A11*, *CD274*, and *CXCL1* were expressed at lower levels in DLBCLs (Figure 2B). Biological processes enriched in DLBCL cMO and iMO included apoptosis, production of ROS, immune response, and phagocytosis (Figure 2C). By flow cytometry, we showed that cMO and iMO from DLBCL displayed a higher expression of CD64 and CCR2 (P < .05), without variation in HLA-DR and CD163 (Figure 2D). Altogether, these results suggested that, in DLBCL, cMO and iMO share a common deregulated inflammatory phenotype.

### DLBCL ncMO are decreased in peripheral blood and exhibit an inflammatory- and tolerogenic- phenotype

We then focused on ncMO in DLBCL samples and found that both subsets of ncMO (ncMO Slan^pos^ and ncMO Slan^neg^) were decreased in DLBCLs when compared to HDs (ncMO Slan^pos^ median at 3.3 x10^6^ cells/L vs 9.1 x10^6^ cells/L [P < .001] and ncMO Slan^neg^ median at 19 x10^6^ cells/L vs 23.7 x10^6^ cells/L [P < .05], respectively) (Figure 3A). No increase of apoptosis was detected in ncMO from DLBCL patients (data not shown). Then, sorted ncMO Slan^pos^ and ncMO Slan^neg^ from DLBCL and HD samples were analyzed by high-throughput Q-PCR (Figure S4). To explore the similarities between ncMO Slan^pos^, ncMO Slan^neg^, iMO, and cMO, we performed a hierarchical clustering on the DLBCL samples. For 6 out of 7 patients, cMO and iMO were separated from ncMO independently of Slan expression (Figure 3B and Figure S5). In addition, ncMO exhibited tolerogenic genes (*PDCD1LG2, IL10, IDO, CD274, AGER, TNFAIP6*) (Figure S5), most of these genes were not expressed in HD ncMO (Figure S6). In DLBCL, cMO and iMO in one hand and ncMO Slan^pos^ and ncMO Slan^neg^ in the other hand shared similar gene expression (Figure 2A and Figure S5), thus we compared the gene expression between ncMO, irrespectively of the Slan status, and both cMO and iMO. DLBCL ncMO were enriched for both inflammatory (*CXCL10, AIM2, IL12A*) and tolerogenic (*PDCD1LG2, IL10, IDO, CD274, AGER*) genes (P < .05, ∣log2FC∣ > 1) compared to cMO and iMO (Figure 3C). Biological processes involved by genes enriched in ncMO from DLBCL patients were growth of tumor, inhibition of cells, and chemotaxis (Figure 3D and Figure S7).

### Tumor conditioned monocytes give rise to ncMO-like prone to migrate in response to CCL3, CCL5, and CXCL12

In order to evaluate how tumor B cells contribute directly to the phenotype of DLBCL monocytes, we cultured monocytes from HD with supernatants from the DLBCL cell lines OCI-Ly3 and OCI-Ly19. After coculture, we analyzed the monocyte phenotype by mass cytometry.^23,25^ After dimension reduction and clustering (Figure 4A), we noticed an increased expression of Slan, CD64, CD163, CD86, CD206, and CD16 and a decreased expression of CCR2 in clusters from tumor-conditioned monocytes when compared to non-conditioned counterparts. We concluded that tumor conditioned monocytes acquired a ncMO-like phenotype. Then we wondered if these cells were prone to migrate into tissue. Tumor-conditioned monocytes demonstrated an increase in *in vitro* migration in response to CCL3 (46.4 and 18.2 fold for OCI-Ly3 and OCI-Ly19, respectively), CCL5 (78.4 and 63.4 fold for OCI-Ly3 and OCI-Ly19, respectively), and CxCL12 (131 and 77.4 fold for OCI-Ly3 and OCI-Ly19, respectively), when compared to non-conditioned monocytes (Figure 4B). Then we compared the phenotype obtained by mass cytometry of tumor-conditioned monocytes (Mo OCI-Ly) to the phenotype of myeloid cells from DLBCL tumors, already published by our group (Flow Repository FR-FCM-Z2CA).^23^ Interestingly, a subset of myeloid cells from DLBCL showed similarities with tumor-conditioned monocytes, in particular with the common high expression of CD64, CD11c, CD32, S100A9, and HLA-DR (Figure 4C).

**Figure 4:**
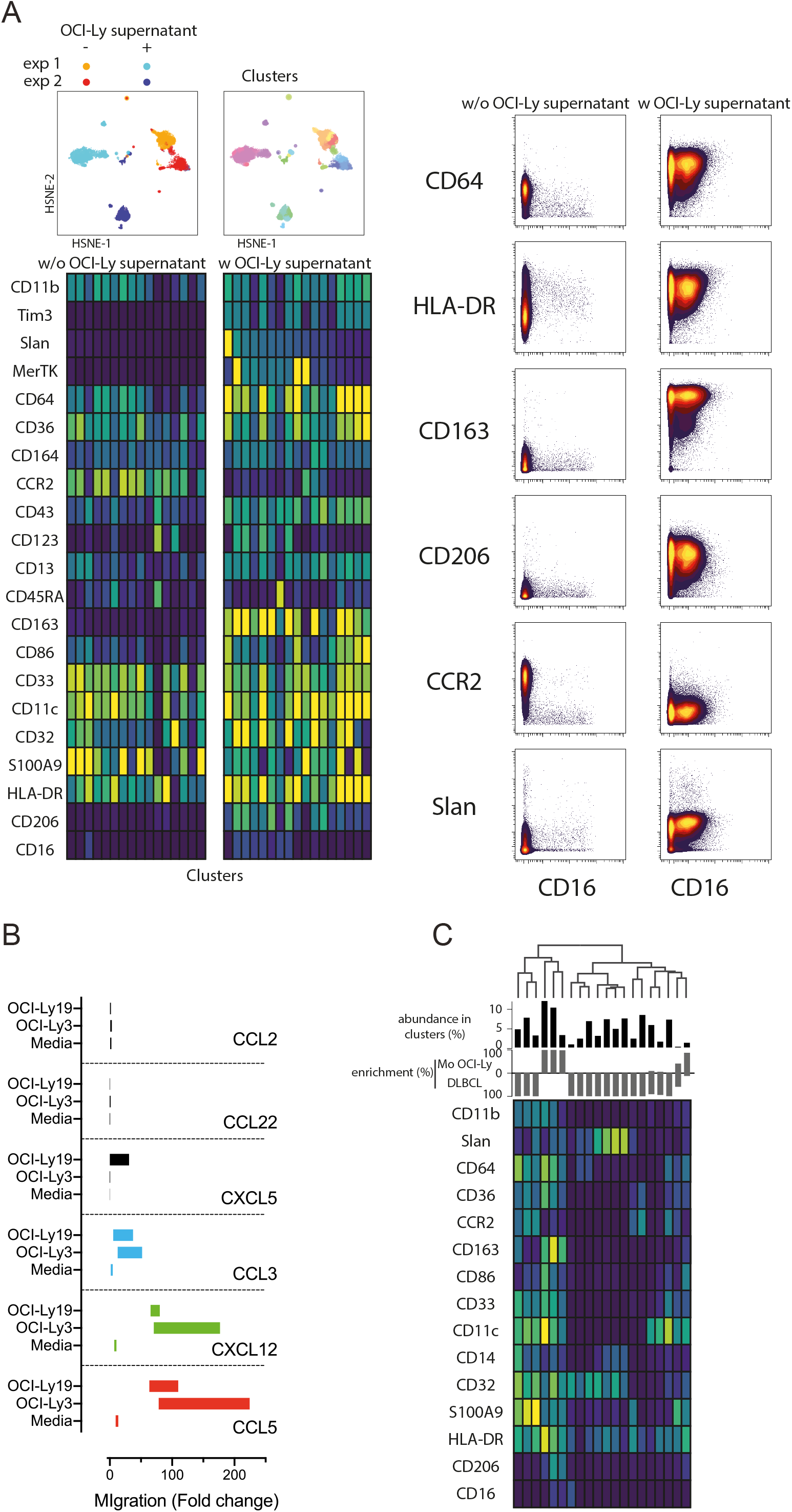
Tumor cells supernatants polarize monocytes with higher migratory abilities. (A) Monocytes from healthy donors were treated with OCI-Ly supernatant (n = 2) or vehicle as control (n = 2). After CyTOF analysis, myeloid cells (12,000 to 30,000) were clustered (n = 33 clusters). Mean marker intensities are shown on a heatmap for each cluster (left). Selected markers are shown for monocytes treated or not with OCI-Ly supernatant (right). (B) Migration assay for HD monocytes cultured or not with OCI-Ly3 and OCI-Ly19 supernatant in response to CCL2, CCL3, CCL5, CCL22, CXCL5, and CXCL12. (C) Phenotype comparison of tumor conditioned monocytes and myeloid cells from DLBCL tumors (n = 7) (FR-FCM-Z2CA, already published by our group).^22^ Mean marker intensities are shown on a heatmap for each cluster. The abundance of clusters is shown as well as the enrichment in tumor-conditioned monocytes (Mo OCI-Ly) and in myeloid cells from DLBCL tumors (DLBCL).

### High level of circulating ncMO is correlated with an adverse prognosis in DLBCL

Then, we evaluated the prognosis value of cMO, iMO, and ncMO in DLBCL. We used i) the proportion of ncMO to other monocytes (ratio ncMO to sum of cMO and iMO) and ii) the absolute count of circulating cMO, iMO, and ncMO. Analysis was performed on 52 patients for which clinical data were available. cMO and iMO were not associated with prognosis (data not shown). By contrast, patients with high proportion of ncMO and high absolute count of circulating ncMO were associated with a lower event-free survival probability (P = .043 and P = .0061, respectively) using thresholds (ratio at 0.06 and ncMO at 20.58 x10^6^ cells/L) defined with the maxstat package (Figure 5A, Figure S8A and Figure S8B).

**Figure 5:**
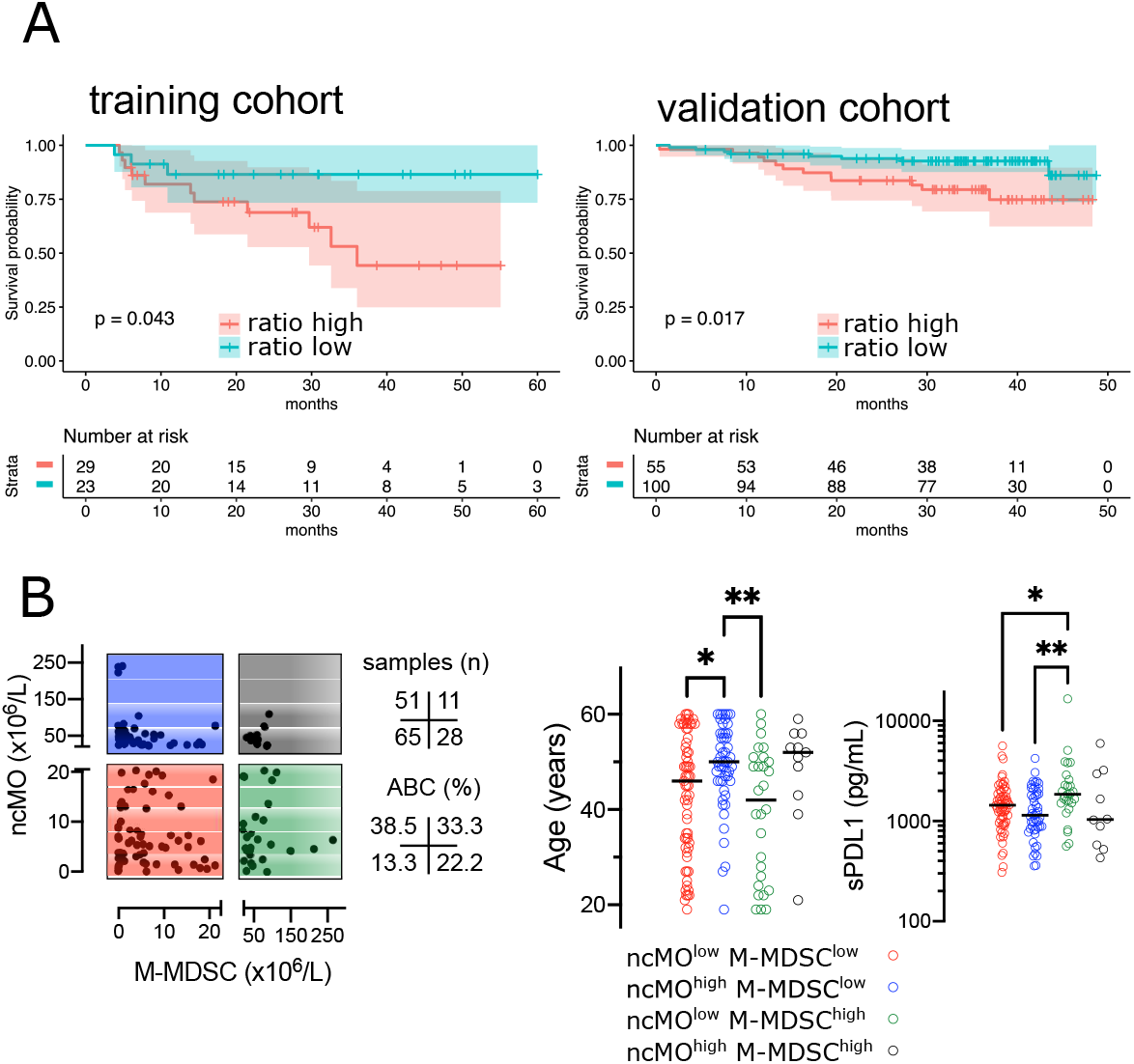
High levels of circulating ncMO is correlated to adverse prognosis in DLBCL. (A) Event-free survival (EFS) in training cohorts (NCT01287923) and overall survival (OS) in validation cohorts (NCT01659099).^24^ Patients were stratified on the ratio of ncMO to other monocytes (cMO and iMO). Threshold was defined on the training cohort using the maxstat package (Figure S8). Survival probability was calculated for both groups with a log-rank test. (B) Absolute count for ncMO (threshold at 20.58 x10^6^ cells/L) and M-MDSC (threshold at 22.51 x10^6^ cells/L)^22^ and distribution of age and soluble PD-L1 (sPD-L1), *P < .05, **P < .01.

To validate the prognosis value of ncMO obtained on this training cohort, we analyzed by flow cytometry the proportion of monocyte subsets in an independent cohort of 155 DLBCL samples from the recently published GAINED trial (NCT01659099).^24^ With the previously calculated thresholds, high proportion of ncMO and high absolute count of ncMO was associated with a lower overall survival (P = .017 and P = .011, respectively) (Figure 5A and Figure S8B). A univariate analysis on the validation cohort showed that Ann Arbor Stage III-IV, ECOG status >1, elevated LDH, PET4 positivity, and increase in circulating ncMO were associated with lower OS (Table S4). In a multivariable analysis Ann Arbor Stage III-IV, PET4 positivity, increase in circulating ncMO remained statistically significant (Table S4).

We previously demonstrated the accumulation of M-MDSC in DLBCL^22^ and since no phenotypic overlap existed between M-MDSC and ncMO (Figure 1D), we wondered if patients’ characteristics were different between M-MDSC^high^ and ncMO^high^ DLBCLs. Both M-MDSC and ncMO were infrequently increased together (11 cases out of 155 [7.1 %]); ncMO were increased alone in 51 cases (32.9 %), and M-MDSC were increased alone in 28 cases (18.1 %) (Figure 5B). Interestingly, ncMO^high^ and M-MDSC^high^ patients corresponded to different types of patients. In particular when compared to ncMO^low^, ncMO^high^ were enriched in ABC DLBCL subtypes (37.5 *vs* 15.9 % [P = .014]) and in older patients (median age at 50 *vs* 46 years [P = .044]). On the other hand, when compared to ncMO^high^, M-MDSC^high^ patients were younger (median age at 42 *vs* 50 years [P = .0027]) and had higher levels of soluble PD-L1 (sPD-L1 at 1849 *vs* 1142 pg/mL [P = .008]) (Figure 5B).

## Discussion

Although the prognostic relevance of total monocyte count has been described in large cohorts of DLBCL in the last decade,^11–15^ few studies evaluated which particular monocyte subset was involved. In a previous work, we have shown an accumulation of M-MDSC (CD14^pos^ HLA-DR^low^) in peripheral blood from DLBCL patients.^22^ Because M-MDSCs were not responsible for the whole increase in monocytes in our cohort, we explored cMO, iMO, and ncMO subsets. We demonstrated an increase in cMOs and iMOs in DLBCL, as in other lymphomas subtypes tested (CLL, MCL, MZL, and FL). In DLBCL, these MO subsets shared an inflammatory phenotype. By contrast, ncMOs were decreased in peripheral blood only in DLBCLs when compared to HDs or other B cell lymphomas. Interestingly, high number of circulating ncMO was an adverse prognosis in 2 independent cohorts of DLBCL patients. Finally, we found that tumor-conditioned monocytes shared a common phenotype with ncMOs and were prone to migrate in response to chemokines.

Surprisingly cMO and iMO from DLBCL shown common deregulated pathways with an enrichment for *FCGR3A, CD36, FCGR1A, CYBB, AIM2, STAT6, FCGR2A, CCR2, NLRC4, S100A8*, and *CD14*. These genes are broadly expressed in cMO in healthy samples^2,5^ and our results suggest that iMO and cMO are tumor-educated and polarized to a common inflammatory phenotype in DLBCL. In our study we found a decrease in both circulating ncMO Slan^neg^ and ncMO Slan^pos^ when compared to HDs, whereas an increase in ncMO Slan^pos^ was previously described in DLBCL.^16^ This discrepancy might be explained by differences in patient characteristics between both studies. In particular patients were older in the study from Verni et coll (63.9 years [range: 31-86] *vs* 50 years [range:18-83]) and at higher grade (clinical stage III-IV at 80.5% *vs* 70% and IPI ≥3 at 55.6% *vs* 40%).^16^ In CLL, an increase of ncMO correlates with high cytogenetic risk (deletion 11q, 17p, or trisomy 12).^30^ In our study, an increase of the proportion of circulating ncMO was a worse prognosis factor in 2 independent cohorts. This was previously suggested on 45 DLBCLs where the decrease of CD16^pos^ monocyte to CD16^neg^ monocyte ratio predicted poor prognosis, however conclusions were limited because iMO and cMO were analyzed conjointly.^17^ ncMO abundance also predicted patient survival of pediatric and adult B acute lymphoblastic leukemia.^31^ Interestingly, in a pre-clinical mouse model of B cell lymphomas, Ly6C^low^ monocytes (corresponding to the ncMO)^32^ accumulated and showed high levels of immunosuppressive genes (*PD-L1, PD-L2, Arg1, IDO1, and CD163*) associated with suppression of T cell proliferation.^33^ In colorectal cancer Ly6C^low^ monocyte mediated immunosuppression by IL-10 production.^34^ Finally, ncMO were increased in gastric cancer.^35^ Conversely, in a lung cancer model, LyC^low^ monocytes recruited NK cells to prevent cancer metastasis.^36^ In DLBCL, we and others focused on total monocyte and on M-MDSC and few attention was given to other monocyte subsets. Interestingly, ncMO and M-MDSC have non-overlapping phenotype regarding HLA-DR expression and these cells infrequently correlated in patients suggesting different mechanism of myelopoiesis dysregulation. Patients that were enriched in circulating MDSCs were younger and presented high amount of sPD-L1, a pejorative marker.^37^ Interestingly, release of PD-L1 was a mechanism of immune suppression suggested in DLBCL.^22^

Beside immunosuppression, gene enriched in ncMO were related to chemotaxis. Circulating ncMO are diminished in DLBCL, on the contrary there were enriched in other B cell lymphomas or solid tumor,^38^ thus we hypothesized that these cells might migrate into tissue to contribute to the tumor-associated macrophage compartment. CCL2, CCL3, CCL5, CCL22, CXCL5, and CXCL12 are involved in monocyte, MDSC, and macrophage recruitment into the TME.^39^ Tumor-conditioned monocyte shown an increased migration in response to CXCL5, CXCL12, CCL3, and CCL5. Consistently, in our previous study, *CXCL5* expression was increased in peripheral blood from DLBCL compared to healthy donors and its expression was related to a worse event-free survival.^22^ CCL3 is also increased in DLBCL when compared to HD and high level correlates with shorter survival.^40,41^

In DLBCL, TAM are heterogenous,^23^ in particular a Slan^pos^ macrophage subset is involved in rituximab mediated antibody dependent cellular cytotoxicity.^16^ In agreement, we found in DLBCL a compartment of cells expressing Slan at high level with CD14, CD32, and HLA-DR. However, DLBCL clusters that correlated with tumor-conditioned monocytes highly expressed CD64, CD36, and S100A9 and thus presented similarities with IFNγ *in-vitro* polarized macrophages.^25^ Few studies compared paired samples from circulating and in situ myeloid cells. In melanoma patient, myeloid cells obtained from the blood, but not from the tumor, were suppressive.^42^ In lung adenocarcinoma, macrophages phenotype detected in tumor were not present in peripheral blood.^43^ Currently, there is no model of lymphoma that allows tracking the myeloid cell from the blood to the tissues. Future studies entailing a prospective collection of paired blood and tumor samples are needed to confirm these observations on ncMO and to put in perspective the myeloid compartment with the T/NK compartment. Also, it would be interested to test the prognosis value in cohort of DLBCL treated with other immunotherapies and correlate with responders vs non-responders.

Our study as some limitations, in particular the lack of extensive functional studies due to the low number of circulating ncMO in DLBCL samples precluding large cell sorting. Taken together, our results show that ncMO are involved in the DLBCL physiopathology and impact the prognosis of the disease. Given the current and our previous data, we propose that cMO and iMO are reflecting the inflammatory status in DLBCL, whereas M-MDSC are responsible of a systemic suppressive response, and ncMO are involved in suppressive response and migration to tissue.

## Acknowledgements

This work was supported by a fellowship from the Nuovo-Soldati Foundation (Switzerland) (M.R.), from the Ligue contre le Cancer (M.R.), from the COmite de la REcherche Clinique et Translationnelle, CHU of Rennes (F.L.), from the Association pour le Développement de l’Hématologie Oncologie (F.L.), from the National Institute of Cancer (INCa Recherche Translationnelle 2010) (T.F.), and from the Groupe Ouest-Est des Leucémies et des Autres Maladies du Sang (GOELAMS) (T.F.). We are indebted to the clinicians of the BREHAT network and to the French Blood Bank (EFS) of Rennes for providing samples. The authors acknowledge the Centre de Ressources Biologiques (CRB-santé) of Rennes (BB-0033-00056) for managing samples.

## Authorship contributions

S.L.Ga., F.L., A.M., C.M., and I.A. designed and performed experiments, analyzed data; J.M.I., D.R., J.F., and T.J.M. analyzed data; C.P., K.B., G.D., G.C., P.G.,

S.L.Go., R.O.C., R.H., and T.L. provided samples; T.F. and K.T. raised the funds and analyzed data; M.R. designed and supervised research, analyzed data, and wrote the paper. All authors revised the manuscript.

