## Supplemental Tables and Figures for "Nonclassical monocytes are prone to migrate into tumor in diffuse large B-cell lymphoma"

**Supplemental figure legends**

**Figure S1: Gating strategy**

**Figure S2 (related to Figure 1): All monocyte subsets are increased in B-cell lymphomas**

A- Monocyte subsets counts in peripheral blood from HD (n = 6), follicular lymphomas (n = 9), mantle cell lymphomas (MCL, n = 9), chronic lymphocytic leukemias (CLL, n = 11), and marginal zone lymphomas (SMZL, n = 10). On these patients, flow cytometry was performed on cryopreserved cells. B- Ratio between MO (classical and intermediate) and ncMO in B-cell lymphomas.

**Figure S3 (related to Figure 2): Q-PCR analysis, hierarchical clustering for cMO and iMO from DLBCL and HD patients**

cMO and iMO were sorted from 7 DLBCL and 4 HD before analysis by high-throughput Q-PCR for genes listed in Table S3. Pearson's correlation and complete linkage were employed.

**Figure S4 (related to Figure 3): Q-PCR analysis, hierarchical clustering for ncMO Slan$^{pos}$ and ncMO Slan$^{neg}$ from DLBCL and HD patients**

ncMO Slan$^{pos}$ and ncMO Slan$^{neg}$ were sorted from 7 DLBCL and 4 HD before analysis by high-throughput Q-PCR for genes listed in Table S3. Pearson's correlation and complete linkage were employed.

**Figure S5 (related to Figure 3): Q-PCR analysis, hierarchical clustering for cMO, iMO, ncMO Slan$^{pos}$ and ncMO Slan$^{neg}$ from DLBCL patients**

cMO, iMO, ncMO Slan$^{pos}$, and ncMO Slan$^{neg}$ were sorted from 7 DLBCL before analysis by high-throughput Q-PCR for genes listed in Table S3. Pearson's correlation and complete linkage were employed.

**Figure S6 (related to Figure 3): Q-PCR analysis, hierarchical clustering for cMO, iMO, ncMO Slan$^{pos}$ and ncMO Slan$^{neg}$ from DLBCL and HD patients**

cMO, iMO, ncMO Slan$^{pos}$, and ncMO Slan$^{neg}$ were sorted from 7 DLBCL and 4 HD before analysis by high-throughput Q-PCR for genes listed in Table S3. Pearson's correlation and complete linkage were employed.

**Figure S7 (related to Figure 3): Transcripts differentially expressed in ncMO and top biological process involved**

Transcripts differentially expressed ($P < .05$; $|log2FC| > 1$) between DLBCLs (n=7) and HDs (n=4), for ncMO Slan$^{pos}$ and ncMO Slan$^{neg}$ (Top). Predicted top 5 biological processes increased for ncMO Slan$^{pos}$ and ncMO Slan$^{neg}$ from DLBCL when compared to HD samples (Ingenuity Pathway Analysis, z-score > 2.5, ranked by p-value) (Bottom).

**Figure S8 (related to Figure 5): High levels of circulating ncMO is an adverse event in DLBCL.**

(A) Event-free survival (EFS) in training cohorts (NCT01287923) Threshold of the ratio of ncMO to other monocytes parameter was defined on the training cohort using the maxstat package. (B) Event-free survival (EFS) in training cohorts (NCT01287923) and overall survival (OS) in validation cohorts (NCT01659099). Patients were stratified on the absolute count of circulating ncMO. Threshold was defined on the training cohort using the maxstat package. Survival probability was calculated for both groups with a log-rank test.

**Table S1: Antibodies for fluorescent flow cytometry analysis**

| Target | Fluorochrome | Clone | Company |
|---|---|---|---|
| HLA-DR | PE-CF594 | G46-6 | BD Biosciences |
| CD14 | PC7 | RMO52 | Beckman Coulter |
| CD16 | APC-Alexa700 | 3G8 | Beckman Coulter |
| CD3 | V450 | UCHT1 | BD Biosciences |
| CD335 | BV421 | 9E2/NKp46 | Biolegend |
| CD45 | Krome Orange | J33 | Beckman Coulter |
| Slan | FITC | DD-1 | Miltenyi Biotec |

### Table S2: Antibodies or parameters used for mass cytometry analysis

| Target / Compound | Metal / Parameter | Clone | Company | Staining |
|---|---|---|---|---|
| CD11b | 141Pr | ICRF44 | Biolegend | C |
| CD19 | 142Nd | HIB19 | Fluidigm | - |
| CD366 Tim3 | 143Nd | F38-2E2 | Biolegend | C |
| Slan-FITC / anti-FITC | 144Nd | DD-1 | Miltenyi Biotech / Fluidigm | I |
| MerTK-PE / anti-PE | 145Nd | 125518 | R&D systems / Fluidigm | I |
| CD64 | 146Nd | 10.1 | Fluidigm | - |
| CD36 | 147Sm | 5-271 | Biolegend | C |
| CD164 | 148Nd | 67D2 | Biolegend | C |
| CCR2 | 149Sm | K036C2 | Biolegend | C |
| CD43 | 150Nd | 84-3C1 | Fluidigm | - |
| CD123 | 151Eu | 6H6 | Fluidigm | - |
| CD13 | 152Sm | WM15 | Fluidigm | - |
| CD45RA | 153Eu | HI100 | Fluidigm | - |
| CD163 | 154Sm | GHI/61 | Fluidigm | - |
| CD27 | 155Gd | L128 | Fluidigm | - |
| CD86 | 156Gd | IT2.2 | Fluidigm | - |
| CD33 | 158Gd | WM53 | Fluidigm | - |
| CD11c | 159Tb | Bu15 | Fluidigm | - |
| CD14 | 160Gd | M5E2 | Fluidigm | - |
| CD32 | 161Dy | FUN-2 | Biolegend | C |
| S100A9-APC / anti-APC | 162Dy | MRP-14 | Biolegend / Fluidigm | I/intra |
| HLA-DR | 163Dy | L243 | Biolegend | C |
| CD206 | 164Dy | 3.29B1.10 | Beckman Coulter | C |
| CD16 | 165Ho | 3G8 | Fluidigm | - |
| CD120a | 166Er | 80M2 | Beckman Coulter | C |
| CCR7 | 167Er | G043H7 | Fluidigm | - |
| CD8 | 168Er | SK1 | Fluidigm | - |
| CD25 | 169Tm | 2A3 | Fluidigm | - |
| CD3 | 170Er | SP34-2 | Fluidigm | - |
| CD68 | 171Yb | Y1/82A | Fluidigm | Intra |
| CD9 | 172Yb | SN4 C3-3A2 | Fluidigm | - |
| CD45 | 173Yb | 2D1 | Biolegend | C |
| CD279 | 174Yb | EH12.2H7 | Biolegend | C |
| CD274 | 175Yb | 29E.2A3 | Fluidigm | - |
| CD127 | 176Yb | A019D5 | Fluidigm | - |
| Iridium | 191Ir | - | Fluidigm | - |
| Iridium | 193Ir | - | Fluidigm | - |
| Cisplatin | 195Pt | - | Enzo Life Sciences | - |
| - | Cell length | - | - | - |

Staining: C: Custom conjugate; I: Indirect staining; intra: intrastaining

**Table S3: List of genes evaluated by high-throughput Q-PCR on monocyte subsets**

| | | | |
|---|---|---|---|
| *ADAM17* | *CD64* | *IL12A* | *RELB* |
| *AIM2* | *CD68* | *IL17R* | *S100A12* |
| *BclxL* | *CD80* | *IL6* | *S100A8* |
| *CEBPb* | *CD86* | *IL6R* | *S100A9* |
| *C5aR* | *CTLA4* | *LAG3* | *SLC7A11* |
| *caspase1* | *CXCL1* | *MMP9* | *STAT1* |
| *CCR2* | *CXCL10* | *MYD88* | *STAT3* |
| *CCR5* | *CyclinD1* | *NFKBP50* | *STAT6* |
| *CD11b* | *EP24* | *NFKBP52* | *TGFb* |
| *CD137* | *GCSFR* | *NIK* | *TGM2* |
| *CD14* | *Galectin9* | *NLRC4* | *Tim3* |
| *CD16* | *GMCSFR* | *NOX2* | *TLR2* |
| *CD163* | *HIFa* | *OX40* | *TLR4* |
| *CD32* | *HLADR* | *PD1* | *TLR6* |
| *CD33* | *HO1* | *PDL1* | *TNFa* |
| *CD36* | *IDO* | *PDL2* | *TNFAIP6* |
| *CD38* | *IL4R* | *RAGE* | *TNFR2* |
| *CD40* | *IL10* | *RELA* | |

### Table S4: Factors influencing overall survival in validation cohort

| Risk factors | | N (%) | Univariate analysis HR | Univariate analysis P | Multivariate analysis HR | Multivariate analysis P |
|---|---|---|---|---|---|---|
| **Age (years)** | >50 | 59 (38.1%) | 1.096 | 0.84 | | |
| | ≤50 | 96 (61.9%) | | | | |
| **Gender** | Male | 86 (55.5%) | 1.027 | 0.952 | | |
| | Female | 69 (44.5%) | | | | |
| **ECOG** | ≥2 | 18 (11.6%) | **3.61** | **0.008** | 1.567 | 0.440 |
| | 0-1 | 137 (88.4%) | | | | |
| **Ann Arbor stage** | III-IV | 130 (83.9%) | **7.52** | **0.006** | **8.42** | **0.002** |
| | I-II | 25 (16.1%) | | | | |
| **LDH** | Elevated | 52 (34.4%) | **2.852** | **0.025** | 2.432 | 0.131 |
| | Normal | 99 (65.6%) | | | | |
| **aaIPI** | 2-3 | 87 (56.1%) | 1.447 | 0.431 | | |
| | 0-1 | 68 (43.9%) | | | | |
| **Bulk** | ≥10 cm | 47 (30.3%) | 1.283 | 0.595 | | |
| | <10 cm | 108 (69.7%) | | | | |
| **COO** | ABC | 83 (74.8%) | 1.704 | 0.302 | | |
| | GCB | 28 (25.2%) | | | | |
| **BCL2** | ≥70 % | 87 (67.4%) | 0.5 | 0.283 | | |
| | <70 % | 42 (32.6%) | | | | |
| **MYC** | ≥40 % | 53 (46.5%) | 1.172 | 0.767 | | |
| | <40% | 61 (53.6%) | | | | |
| **DE MYC/BCL2** | Yes | 45 (29.8%) | 0.847 | 0.75 | | |
| | No | 106 (70.2%) | | | | |
| **Treatment arm** | Obinutuzumab | 76 (49.4%) | 1.089 | 0.853 | | |
| | Rituximab | 78 (50.6%) | | | | |
| **Chemotherapy** | CHOP | 77 (50%) | 0.881 | 0.784 | | |
| | ACVBP | 77 (50%) | | | | |
| **PET2/PET4** | PET4+ | 26 (19%) | **3.744** | **0.013** | **2.943** | **0.03** |
| | PET2- or PET2+/PET4- | 111 (81%) | | | | |
| **ncMO ($x10^6$/L)** | ≥20.58 | 62 (40%) | **3.135** | **0.015** | **3.362** | **0.047** |
| | <20.58 | 93 (60%) | | | | |

ECOG: Eastern Cooperative Oncology Group scale; LDH: lactate deshydrogenase; aaIPI: age adjusted International prognostic index; COO: cell of origin; DE: double expressor; PET2: PET after cycle 2; PET4: PET after cycle 4; ncMO: non classical monocyte

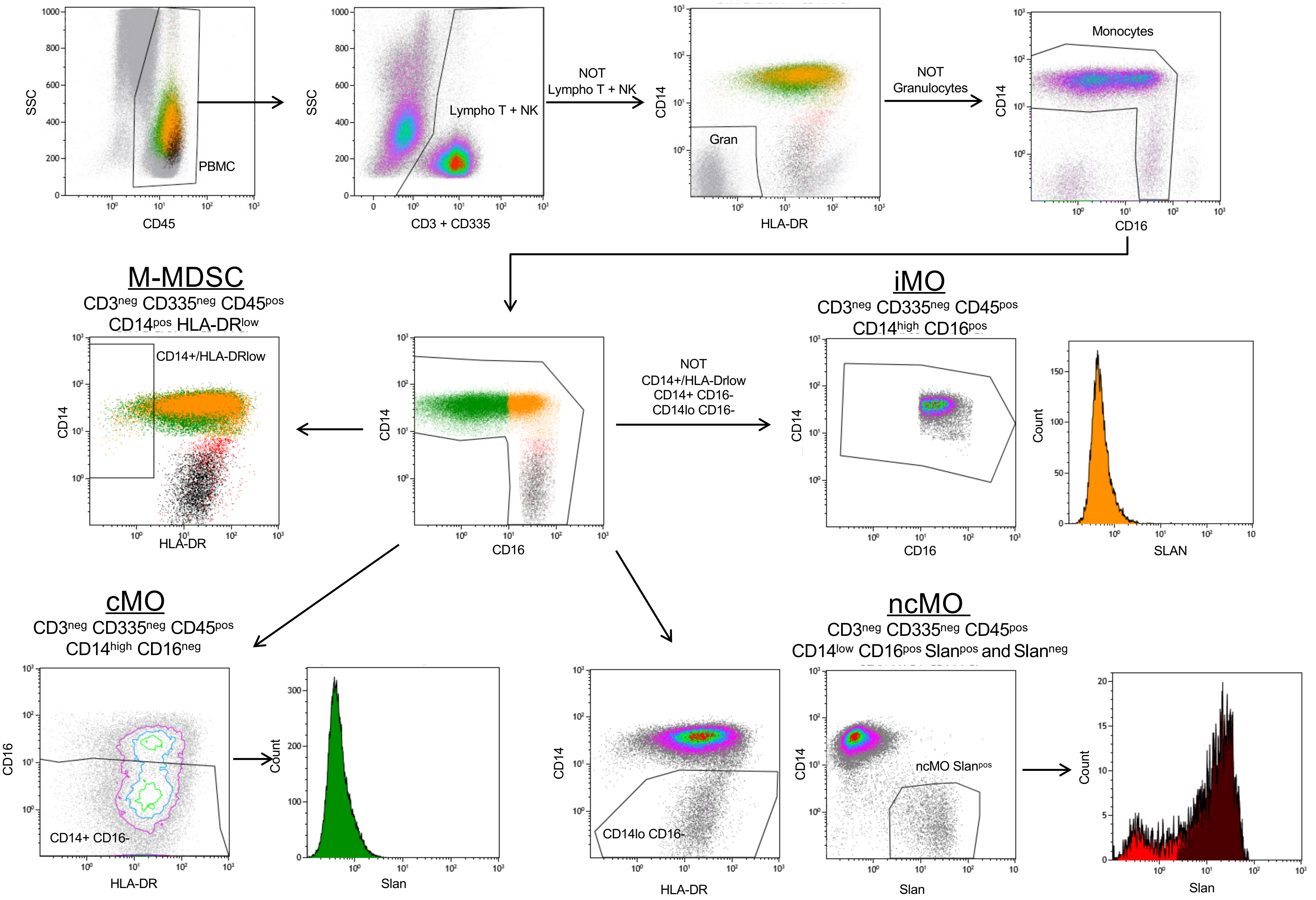

SSC
CD45
PBMC
CD3 + CD335
Lympho T + NK
NOT
Lympho T + NK
CD14
HLA-DR
Gran
NOT
Granulocytes
Monocytes
CD16
M-MDSC
CD3neg CD335neg CD45pos
CD14pos HLA-DRlow
CD14+/HLA-DRlow
NOT
CD14+/HLA-Drlow
CD14+ CD16-
CD14lo CD16-
iMO
CD3neg CD335neg CD45pos
CD14high CD16pos
Count
SLAN
cMO
CD3neg CD335neg CD45pos
CD14high CD16neg
CD14+ CD16-
Slan
ncMO
CD3neg CD335neg CD45pos
CD14low CD16pos Slanpos and Slanneg
CD14lo CD16-
ncMO Slanpos

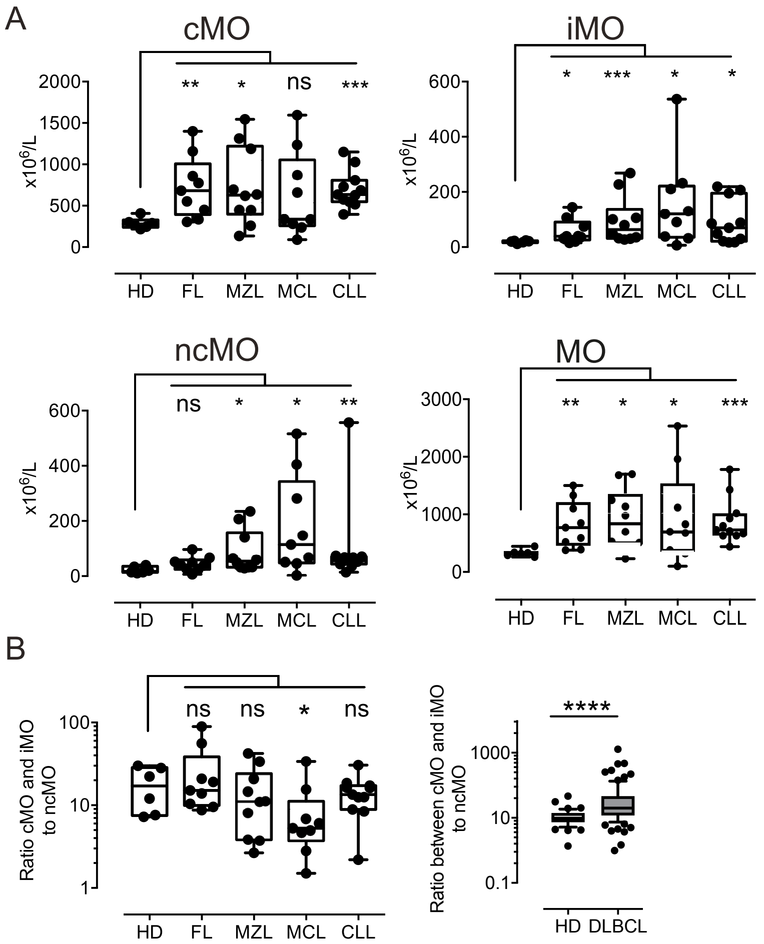
A
cMO
iMO
ncMO
MO
x10^6/L
HD
FL
MZL
MCL
CLL
ns
**
*
***
B
Ratio cMO and iMO to ncMO
Ratio between cMO and iMO to ncMO
****
DLBCL

Figure S2

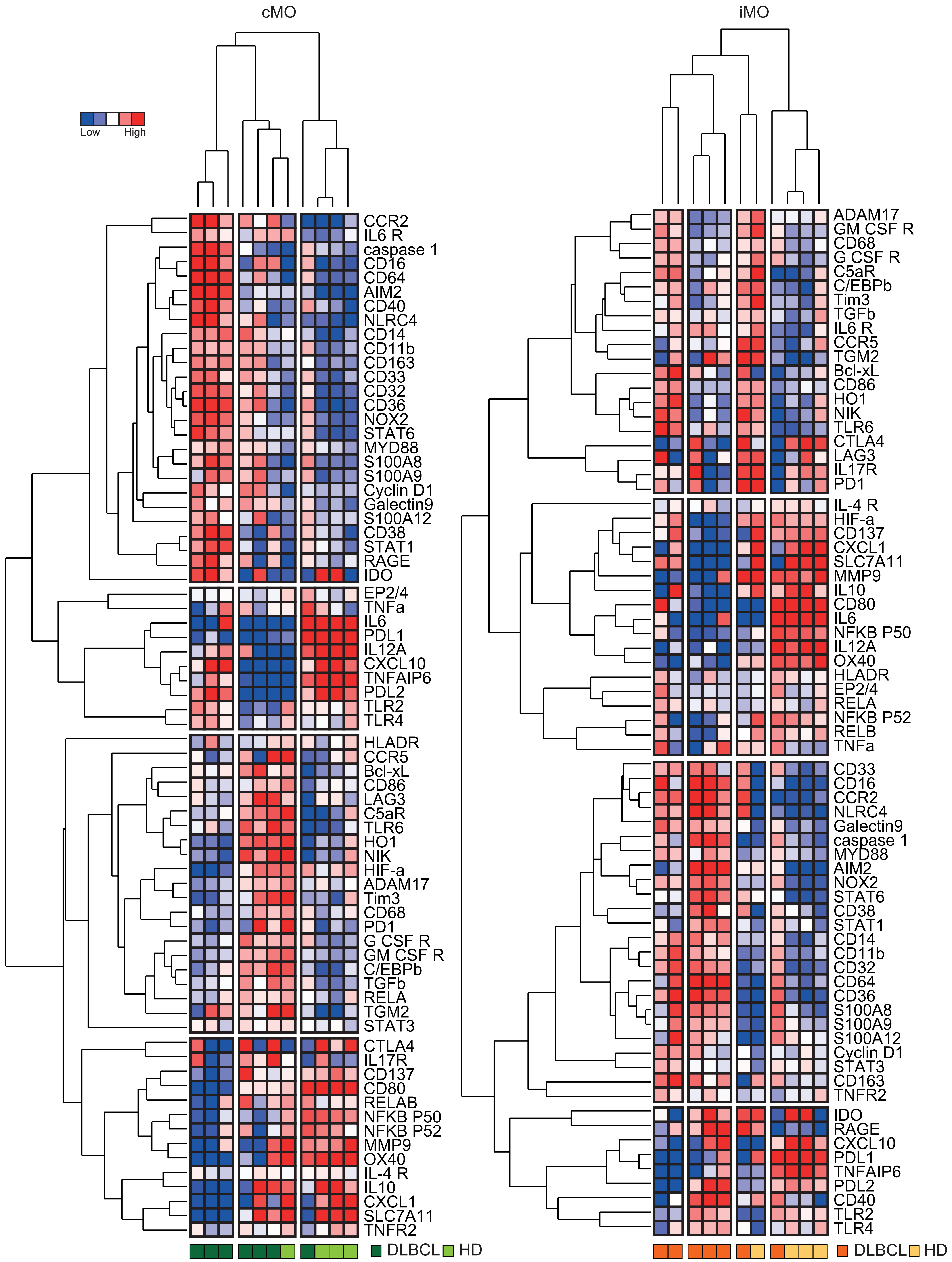
cMO
iMO
Low
High
CCR2
IL6 R
caspase 1
CD16
CD64
AIM2
CD40
NLRC4
CD14
CD11b
CD163
CD33
CD32
CD36
NOX2
STAT6
MYD88
S100A8
S100A9
Cyclin D1
Galectin9
S100A12
CD38
STAT1
RAGE
IDO
EP2/4
TNFa
IL6
PDL1
IL12A
CXCL10
TNFAIP6
PDL2
TLR2
TLR4
HLADR
CCR5
Bcl-xL
CD86
LAG3
C5aR
TLR6
HO1
NIK
HIF-a
ADAM17
Tim3
CD68
PD1
G CSF R
GM CSF R
C/EBPb
TGFb
RELA
TGM2
STAT3
CTLA4
IL17R
CD137
CD80
RELAB
NFKB P50
NFKB P52
MMP9
OX40
IL-4 R
IL10
CXCL1
SLC7A11
TNFR2
DLBCL
HD
ADAM17
GM CSF R
CD68
G CSF R
C5aR
C/EBPb
Tim3
TGFb
IL6 R
CCR5
TGM2
Bcl-xL
CD86
HO1
NIK
TLR6
CTLA4
LAG3
IL17R
PD1
IL-4 R
HIF-a
CD137
CXCL1
SLC7A11
MMP9
IL10
CD80
IL6
NFKB P50
IL12A
OX40
HLADR
EP2/4
RELA
NFKB P52
RELB
TNFa
CD33
CD16
CCR2
NLRC4
Galectin9
caspase 1
MYD88
AIM2
NOX2
STAT6
CD38
STAT1
CD14
CD11b
CD32
CD64
CD36
S100A8
S100A9
S100A12
Cyclin D1
STAT3
CD163
TNFR2
IDO
RAGE
CXCL10
PDL1
TNFAIP6
PDL2
CD40
TLR2
TLR4
DLBCL
HD

Figure S3

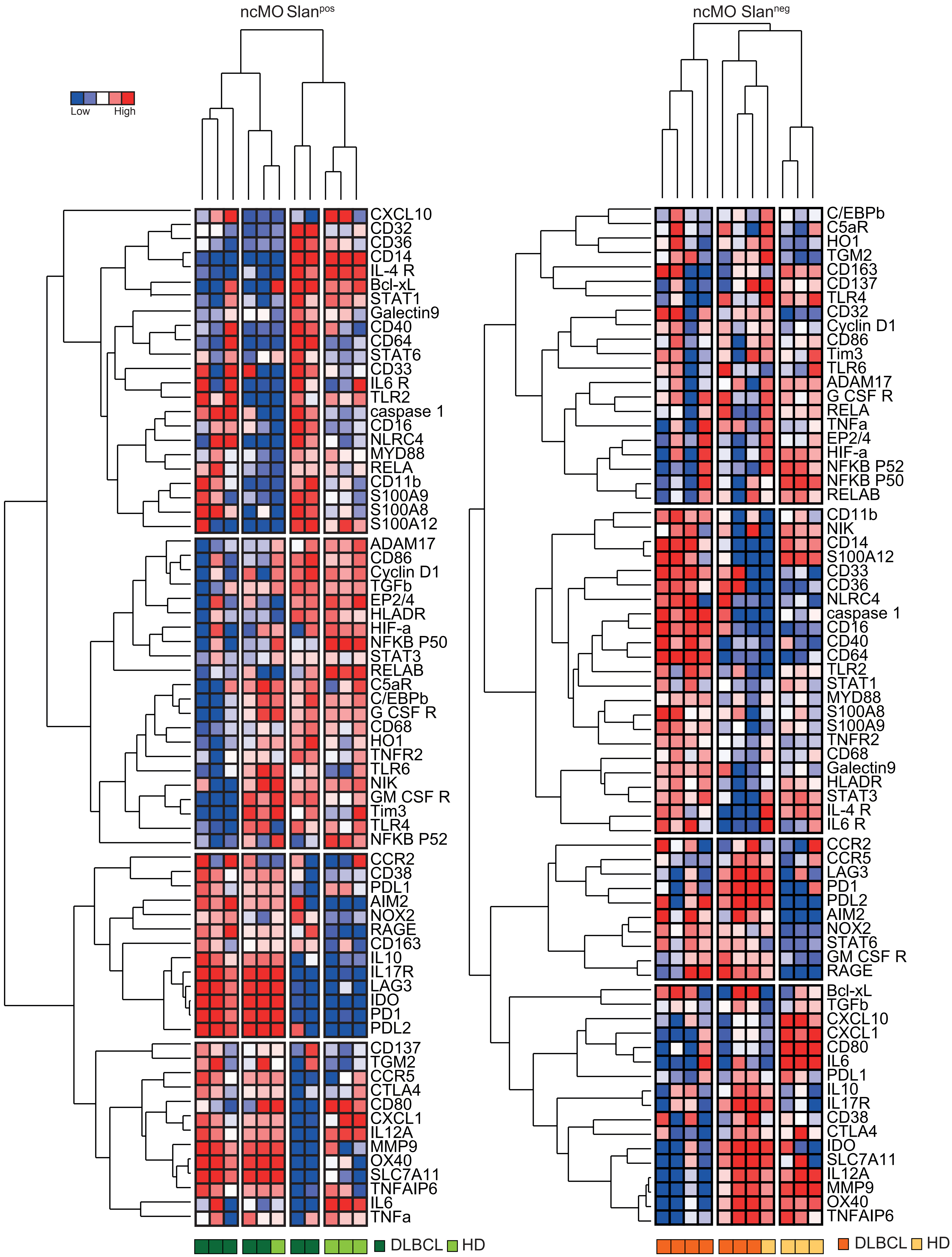

ncMO Slan$^{pos}$
ncMO Slan$^{neg}$
Low
High
CXCL10
CD32
CD36
CD14
IL-4 R
Bcl-xL
STAT1
Galectin9
CD40
CD64
STAT6
CD33
IL6 R
TLR2
caspase 1
CD16
NLRC4
MYD88
RELA
CD11b
S100A9
S100A8
S100A12
ADAM17
CD86
Cyclin D1
TGFb
EP2/4
HLADR
HIF-a
NFKB P50
STAT3
RELAB
C5aR
C/EBPb
G CSF R
CD68
HO1
TNFR2
TLR6
NIK
GM CSF R
Tim3
TLR4
NFKB P52
CCR2
CD38
PDL1
AIM2
NOX2
RAGE
CD163
IL10
IL17R
LAG3
IDO
PD1
PDL2
CD137
TGM2
CCR5
CTLA4
CD80
CXCL1
IL12A
MMP9
OX40
SLC7A11
TNFAIP6
IL6
TNFa
DLBCL
HD
C/EBPb
C5aR
HO1
TGM2
CD163
CD137
TLR4
CD32
Cyclin D1
CD86
Tim3
TLR6
ADAM17
G CSF R
RELA
TNFa
EP2/4
HIF-a
NFKB P52
NFKB P50
RELAB
CD11b
NIK
CD14
S100A12
CD33
CD36
NLRC4
caspase 1
CD16
CD40
CD64
TLR2
STAT1
MYD88
S100A8
S100A9
TNFR2
CD68
Galectin9
HLADR
STAT3
IL-4 R
IL6 R
CCR2
CCR5
LAG3
PD1
PDL2
AIM2
NOX2
STAT6
GM CSF R
RAGE
Bcl-xL
TGFb
CXCL10
CXCL1
CD80
IL6
PDL1
IL10
IL17R
CD38
CTLA4
IDO
SLC7A11
IL12A
MMP9
OX40
TNFAIP6
DLBCL
HD


Figure S4

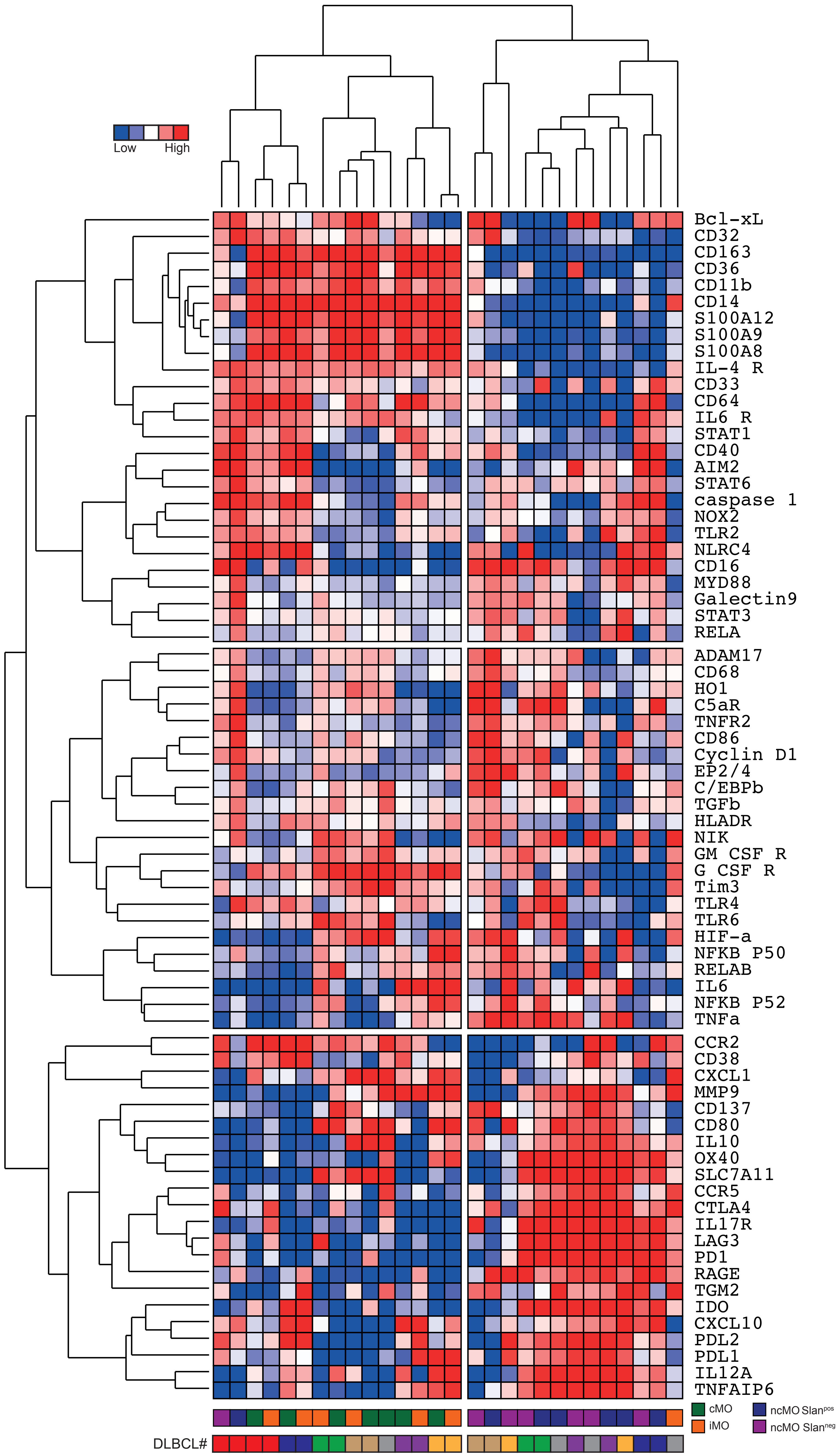
Low
High
Bcl-xL
CD32
CD163
CD36
CD11b
CD14
S100A12
S100A9
S100A8
IL-4 R
CD33
CD64
IL6 R
STAT1
CD40
AIM2
STAT6
caspase 1
NOX2
TLR2
NLRC4
CD16
MYD88
Galectin9
STAT3
RELA
ADAM17
CD68
HO1
C5aR
TNFR2
CD86
Cyclin D1
EP2/4
C/EBPb
TGFb
HLADR
NIK
GM CSF R
G CSF R
Tim3
TLR4
TLR6
HIF-a
NFKB P50
RELAB
IL6
NFKB P52
TNFa
CCR2
CD38
CXCL1
MMP9
CD137
CD80
IL10
OX40
SLC7A11
CCR5
CTLA4
IL17R
LAG3
PD1
RAGE
TGM2
IDO
CXCL10
PDL2
PDL1
IL12A
TNFAIP6
cMO
iMO
ncMO Slanpos
ncMO Slanneg
DLBCL#

Figure S5

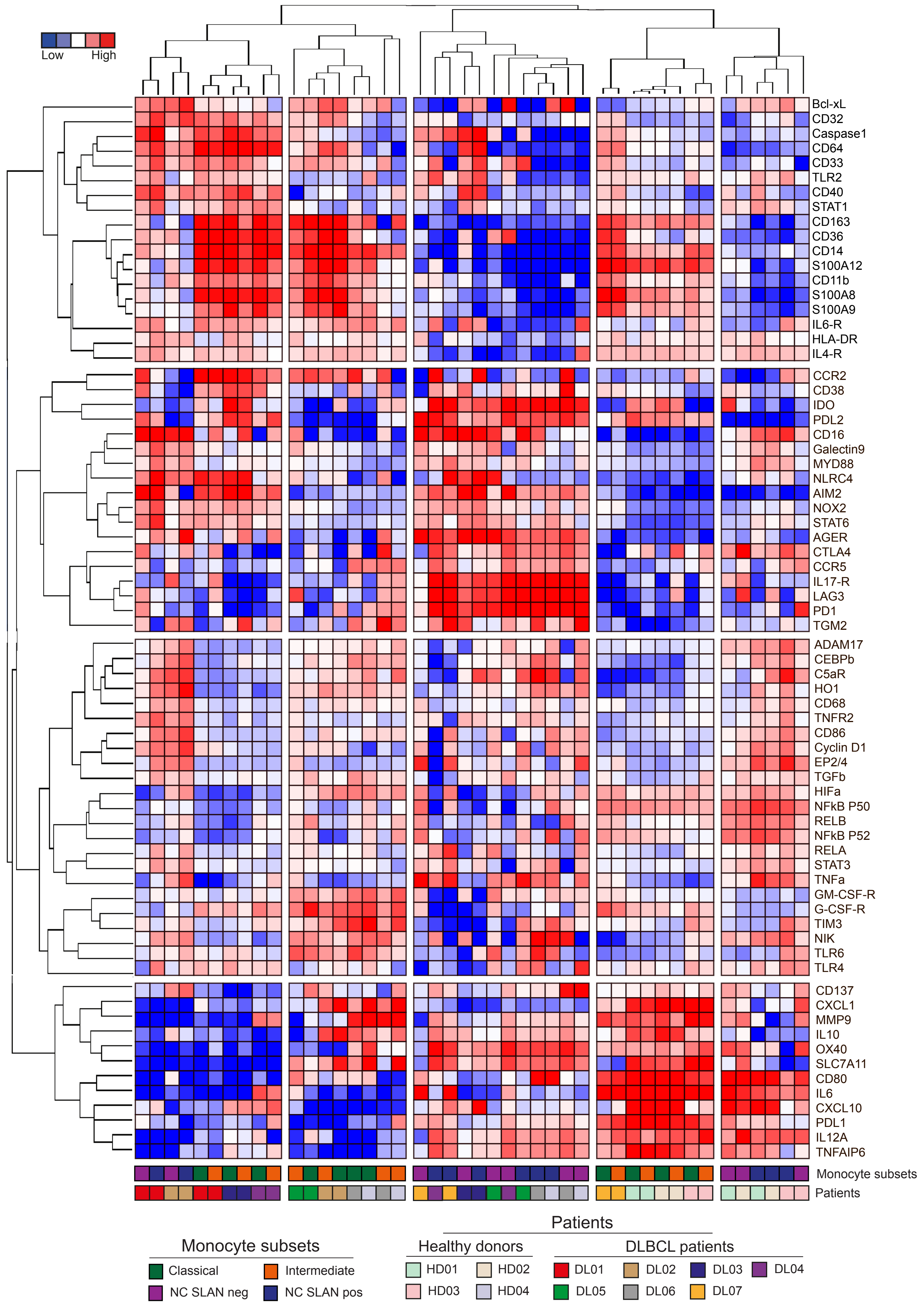

Low
High
Bcl-xL
CD32
Caspase1
CD64
CD33
TLR2
CD40
STAT1
CD163
CD36
CD14
S100A12
CD11b
S100A8
S100A9
IL6-R
HLA-DR
IL4-R
CCR2
CD38
IDO
PDL2
CD16
Galectin9
MYD88
NLRC4
AIM2
NOX2
STAT6
AGER
CTLA4
CCR5
IL17-R
LAG3
PD1
TGM2
ADAM17
CEBPb
C5aR
HO1
CD68
TNFR2
CD86
Cyclin D1
EP2/4
TGFb
HIFa
NFkB P50
RELB
NFkB P52
RELA
STAT3
TNFa
GM-CSF-R
G-CSF-R
TIM3
NIK
TLR6
TLR4
CD137
CXCL1
MMP9
IL10
OX40
SLC7A11
CD80
IL6
CXCL10
PDL1
IL12A
TNFAIP6
Monocyte subsets
Patients
Patients
Monocyte subsets
Classical
Intermediate
NC SLAN neg
NC SLAN pos
Healthy donors
HD01
HD02
HD03
HD04
DLBCL patients
DL01
DL02
DL03
DL04
DL05
DL06
DL07


Figure S6

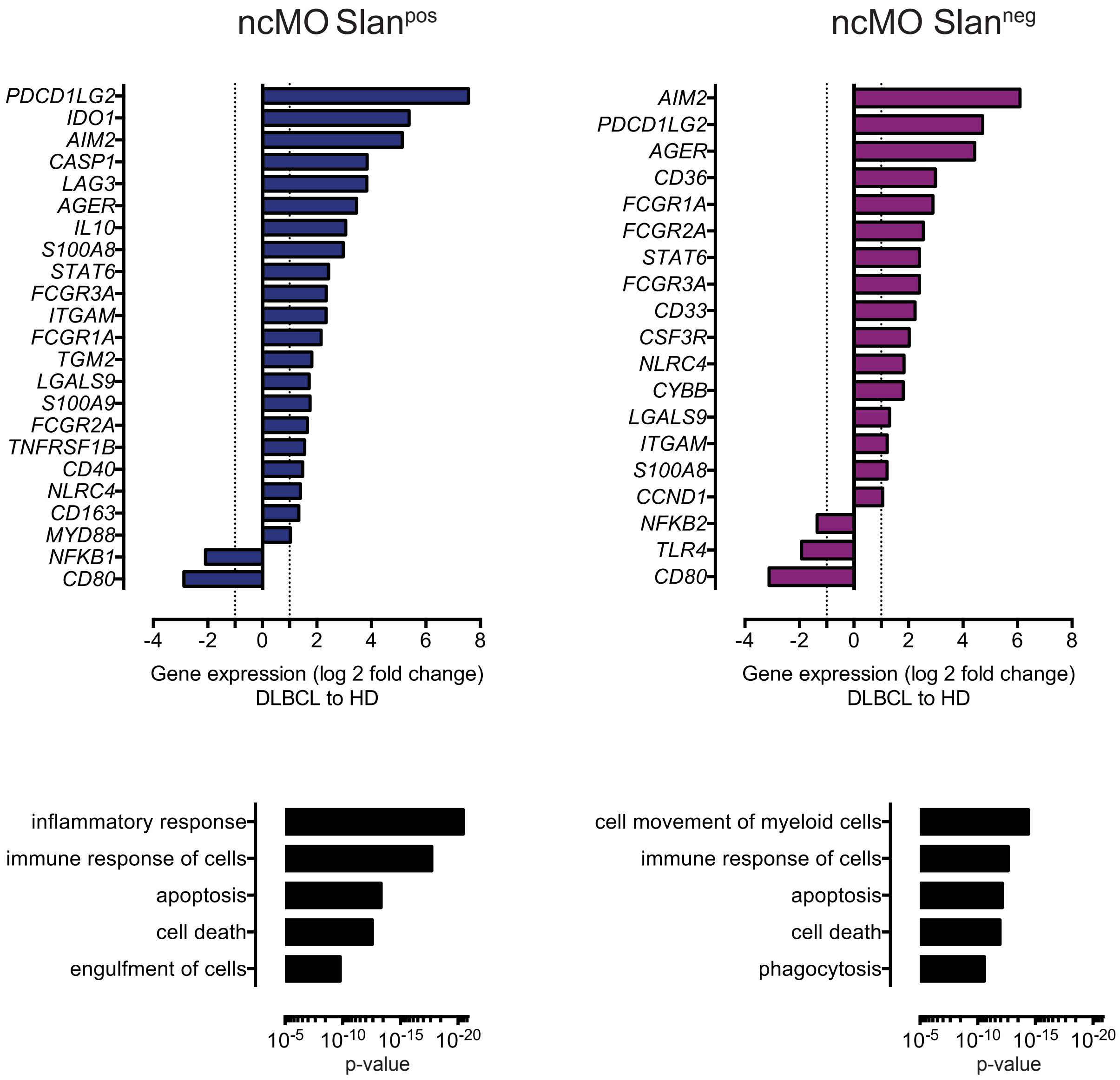
ncMO Slanpos
PDCD1LG2
IDO1
AIM2
CASP1
LAG3
AGER
IL10
S100A8
STAT6
FCGR3A
ITGAM
FCGR1A
TGM2
LGALS9
S100A9
FCGR2A
TNFRSF1B
CD40
NLRC4
CD163
MYD88
NFKB1
CD80
-4 -2 0 2 4 6 8
Gene expression (log 2 fold change)
DLBCL to HD
ncMO Slanneg
AIM2
PDCD1LG2
AGER
CD36
FCGR1A
FCGR2A
STAT6
FCGR3A
CD33
CSF3R
NLRC4
CYBB
LGALS9
ITGAM
S100A8
CCND1
NFKB2
TLR4
CD80
-4 -2 0 2 4 6 8
Gene expression (log 2 fold change)
DLBCL to HD
inflammatory response
immune response of cells
apoptosis
cell death
engulfment of cells
10-5 10-10 10-15 10-20
p-value
cell movement of myeloid cells
immune response of cells
apoptosis
cell death
phagocytosis
10-5 10-10 10-15 10-20
p-value

Figure S7

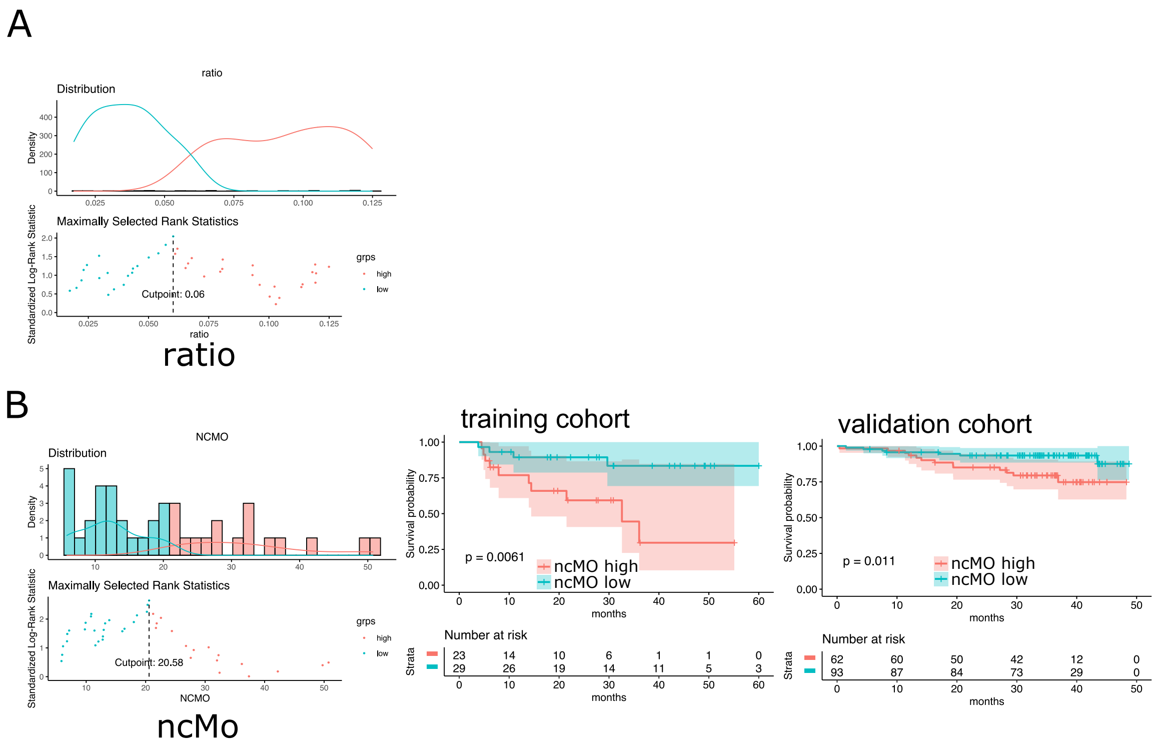
A
ratio
Distribution
Density
Maximally Selected Rank Statistics
Standardized Log-Rank Statistic
Cutpoint: 0.06
grps
high
low
ratio
B
NCMO
Distribution
Density
Maximally Selected Rank Statistics
Standardized Log-Rank Statistic
Cutpoint: 20.58
NCMO
ncMo
training cohort
Survival probability
p = 0.0061
ncMO high
ncMO low
months
Number at risk
Strata
23 14 10 6 1 1 0
29 26 19 14 11 5 3
validation cohort
Survival probability
p = 0.011
ncMO high
ncMO low
months
Number at risk
Strata
62 60 50 42 12 0
93 87 84 73 29 0

Figure S8
